# Towards comprehensive characterization of CRISPR-linked genes

**DOI:** 10.1101/270033

**Authors:** Sergey A. Shmakov, Kira S. Makarova, Yuri I. Wolf, Konstantin V. Severinov, Eugene V. Koonin

**Affiliations:** Skolkovo Institute of Science and Technology, Skolkovo, 143025, Russia; National Center for Biotechnology Information, National Library of Medicine, Bethesda, MD 20894, USA; Waksman Institute for Microbiology Rutgers, The State University of New Jersey, Piscataway, NJ 08854, USA; Institute of Molecular Genetics, Russian Academy of Sciences, Moscow, 123182, Russia

## Abstract

The CRISPR-Cas systems of bacterial and archaeal adaptive immunity consist of arrays of direct repeats separated by unique spacers and multiple CRISPR-associated (cas) genes encoding proteins that mediate the adaptation, CRISPR RNA maturation and interference stages of the CRISPR response. In addition to the relatively small set of core *cas* genes that are typically present in all representatives of each (sub)type of CRISPR-Cas systems and are essential for the defense function, numerous genes occur in CRISPR-cas loci only sporadically. Some of these have been shown to perform various ancillary roles in CRISPR response whereas the functional relevance of many others, if any, remains obscure. We developed a computational strategy for systematically detecting genes that are likely to be functionally linked to CRISPR-Cas systems. The approach is based on a “CRISPRicity” metric that measures the strength of CRISPR association for all protein-coding genes from sequenced bacterial and archaeal genomes. Uncharacterized genes with CRISPRicity values comparable to those of known *cas* genes are considered candidate CRISPR-ancillary genes, and we describe additional criteria to identify functionally relevant genes in the candidate set. About 80 genes that were not previously reported to be associated with CRISPR-Cas were identified as probable CRISPR-ancillary genes. A substantial majority of these genes reside in type III CRISPR-cas loci which implies exceptional functional versatility of type III systems. Numerous candidate CRISPR-ancillary genes encode integral membrane proteins suggestive of tight membrane connections of type III CRISPR-Cas whereas many other candidates are proteins implicated in various signal transduction pathways. These predictions provide ample material for improving annotation of CRISPR-cas loci and experimental characterization of previously unsuspected aspects of CRISPR-Cas functionality.

**SIGNIFICANCE:** The CRISPR-Cas systems that mediate adaptive immunity in bacteria and archaea encompass a small set of core *cas* genes that are essential in a broad range of CRISPR-Cas systems. However, a much greater number of genes only sporadically co-occur with CRISPR-Cas, and for most of these, involvement in CRISPR-Cas functions has not been demonstrated. We developed a computational strategy that provides for systematic identification of CRISPR-linked proteins and prediction of their functional association with CRISPR-Cas systems. About 80 previously undetected, putative CRISPR-accessory proteins were identified. A large fraction of these proteins are predicted to be membrane-associated revealing an unknown side of CRISPR biology.

## Introduction

Driven largely by the exceptional recent success of Cas9, Cas12 and Cas13 RNA-guided nucleases as the new generation of genome and transcriptome editing tools, comparative genomics, structures, biochemical activities and biological functions of CRISPR-Cas systems and individual Cas proteins have been studied in exquisite detail (1–5). The CRISPR-Cas immune response is conventionally described in terms of three distinct stages: 1) adaptation, 2) expression and CRISPR (cr) RNA maturation, and 3) interference. At the adaptation stage, a distinct complex of Cas proteins binds to a target DNA, migrates along that molecule and, typically after encountering a short (2 to 4 bp) motif known as PAM (Protospacer-Adjacent Motif), excises a portion of the target DNA (protospacer), and inserts it into the CRISPR array between two repeats (most often, at the beginning of the array, downstream of the leader sequence) as a spacer (6, 7). The adaptation process creates immune memory, i.e. “vaccinates” a bacterium or archaeon against subsequent infection with the memorized agent. At the expression-maturation stage, the CRISPR array is transcribed into a single, long transcript, the pre-cr(CRISPR)RNA, that is then processed into mature crRNAs, each consisting of a spacer and a portion of an adjacent repeat, by a distinct complex of Cas proteins or a single, large Cas protein, or an external, non-Cas RNase (8). At the final, interference stage, the crRNA that typically remains bound to the processing complex is employed as the guide to recognize the protospacer or a closely similar sequence in an invading genome of a virus or plasmid that is then cleaved and inactivated by a Cas nuclease (s) (9, 10).

Under the current classification, the CRISPR-Cas systems are divided into two classes which radically differ with respect to the composition and structure of the effector modules that are responsible for the interference and, in most of the CRISPR-Cas types, also the processing stages (11, 12). In Class 1 systems, which include types I, III and IV, the effector module is a complex of several Cas proteins which perform a tightly coordinated sequence of reaction, from pre-crRNA processing to target cleavage. In Class 2 systems, including types II, V and VI, all activities of the effector module reside in a single, large multidomain protein such as Cas9 in type II, the programmable endonuclease that is most widely used for genome editing applications (2, 3, 13).

The biochemical activities and biological functions of the 13 families of core Cas proteins that are essential for each of the three stages of the CRISPR immune response in different types of CRISPR-Cas have been extensively studied although some notable gaps in knowledge remains (4, 14, 15). Thus, Cas1 and Cas2 form the adaptation complex that is universal to all autonomous CRISPR-Cas systems (many type III loci and a few in other types lack the adaptation module and apparently rely on the adaptation machinery of other CRISPR-Cas systems in the same organism which they recruit in trans). Cas3 is a helicase that typically also contains a nuclease domain and is involved in target cleavage in type I systems. Cas4 is an endonuclease that is required for adaptation in many CRISPR-Cas variants although its exact function remains unknown. Cas5, Cas6, and Cas7 are distantly related members of the so-called RAMP domain superfamily which includes RNases involved in pre-crRNA processing in types I and III (Cas6 and some variants of Cas5), as well as enzymatically inactive RNA-binding proteins that form the backbone of type I and type III effector complexes. Cas8 is the enzymatically inactive large subunit of type I effector complexes. Cas9 is the type II effector nuclease. Cas10 is the large subunit of the type III effector complexes that contains a Palm domain homologous to those of DNA polymerases and nucleotide cyclases and possesses a oligoA synthetase activity. Cas11 is the small subunit of the type I and type III effector complexes. Cas12 is the effector endonuclease of type V. Finally, Cas13 is the effector RNase of type VI.

In addition to the core proteins included in the formal Cas nomenclature, numerous proteins are found in various subsets of CRISPR-Cas systems and are generally thought of as performing accessory, in particular, regulatory functions in the CRISPR response (4, 11, 16). Admittedly, the separation of the proteins encoded in the CRISPR loci into Cas and accessory groups is somewhat arbitrary because some of the “accessory” ones play important roles in the functionality of the system. As a case in point, many type III systems employ an alternative mechanism of adaptation, namely, spacer acquisition from RNA via reverse transcription by a reverse transcriptase (RT) that is encoded in the CRISPR-*cas* locus, often being fused to the Cas1 protein (17, 18). Another striking case of an “accessory” protein turning out to be an essential component of CRISPR-Cas systems is the recently elucidated function of the Csm6 protein that is also common among type III systems and consists of a nucleotide ligand-binding CARF domain fused to a HEPN RNase domain (19). It has been shown in two independent studies that upon target recognition by Csm6-containing type III systems, Cas10 is induced to synthesize oligoA molecules that are bound by the CARF domain of Csm6 resulting in stimulation of the RNase activity of the HEPN domain which then initiates the immune response (20, 21). However, for most of the putative accessory proteins encoded in CRISPR-cas loci, the functions and the very relevance for the CRISPR response remain unknown. The question of functional relevance is far from being moot because defense islands in microbial genomes appear to be “genomic junkyards” that often accommodate functionally irrelevant genes (22).

We sought to develop a computational strategy for systematic identification of proteins that are non-randomly associated with CRISPR-Cas systems and assess their functional relevance. The approach is based on the CRISPRicity metric that measures the strength of CRISPR association for individual genes. Genes with CRISPRicity values similar to those of *cas* genes were considered candidates for previously undetected accessory genes, and the encoded proteins were examined in depth using sensitive methods for domain identification and coevolution analysis. More than 100 genes encoding diverse proteins were predicted to be involved in CRISPR-Cas functions although not connected to CRISPR-Cas previously. A substantial majority of the detected putative CRISPR-accessory proteins are encoded in type III CRISPR-*cas* loci, and many of these are implicated in membrane association of these systems and/or various signaling pathways.

## RESULTS AND DISCUSSION

### CRISPR-linked genes predicted using CRISPRicity

In bacteria and archaea, functionally linked genes are often organized into operons, i.e. arrays of co-directed, co-transcribed and co-translated genes (23–25). The evolutionary dynamics of operons is such that, for a group of genes that are involved in the same pathway or functional system, different microbial genomes often contain partially overlapping operons encoding subsets of proteins involved in the respective process (26). Comparative analysis of gene neighborhoods in multiple genomes often provides for a complete delineation of the components of functional systems. Indeed, precisely this type of analysis resulted in the description of the group of functionally linked proteins that later became known as Cas (27).

We sought to expand the set of proteins that are functionally linked to the CRISPR-mediated immunity by enumerating all genes in the CRISPR neighborhoods and attempting to distinguish functionally relevant genes from spurious ones. To this end, a dedicated computational pipeline was constructed (Figure 1). Briefly, we identified the union of all genes located in the vicinity (±10 kb) of CRISPR arrays, *cas1* genes (representing the CRISPR-Cas adaptation module) or CRISPR effector modules (for details, see Methods and Supporting Information File 1), hereinafter, CRISPR-linked genes. The union of all these genes was taken so as to maximize the likelihood of detection of CRISPR-linked genes because certain CRISPR-*cas* loci might lack one or even two these key elements. All genes in the CRISPR-cas neighborhoods were represented as points in 3 dimensions defined by: 1) CRISPRicity, i.e. the ratio of the number of CRISPR-linked occurrences of the given gene to its total number of occurrences in the analyzed genomes, 2) abundance (total number of occurrences), and 3) distance from the respective genomic anchor (CRISPR array, *cas1* or effector). All the genes in the neighborhoods were classified into 1) CRISPR-associated (including both *cas* genes and previously identified accessory genes), 2) non-CRISPR (i.e. genes with well-characterized functions unrelated to the CRISPR immunity), and 3) unknowns. The counts of the genes in each of the 3 classes were obtained for each voxel in the space of the 3 coordinates defined above, and the ratio of the probability mass of CRISPR-associated genes (*D_c_*) to that of the non-CRISPR genes (*D_n_*) was calculated (hereinafter CRISPR-index; see Methods for details). The “unknown” and “non-CRISPR” genes with *D_c_*/*D_n_* > 2 were considered candidates for previously undetected CRISPR-linked genes. This threshold was chosen to select genes with similar characteristics to genes known to be functionally linked to CRISPR-Cas system. Indeed, the range of CRISPR-index values for the candidate accessory proteins identified by this procedure overlapped that for Cas proteins and functionally characterized accessory proteins which is compatible with their functional relevance of the previously undetected candidates (Figure 2A). Clearly, the CRISPR-index cut-off can be adjusted to search for stricter or looser associations.

**Figure 1.**
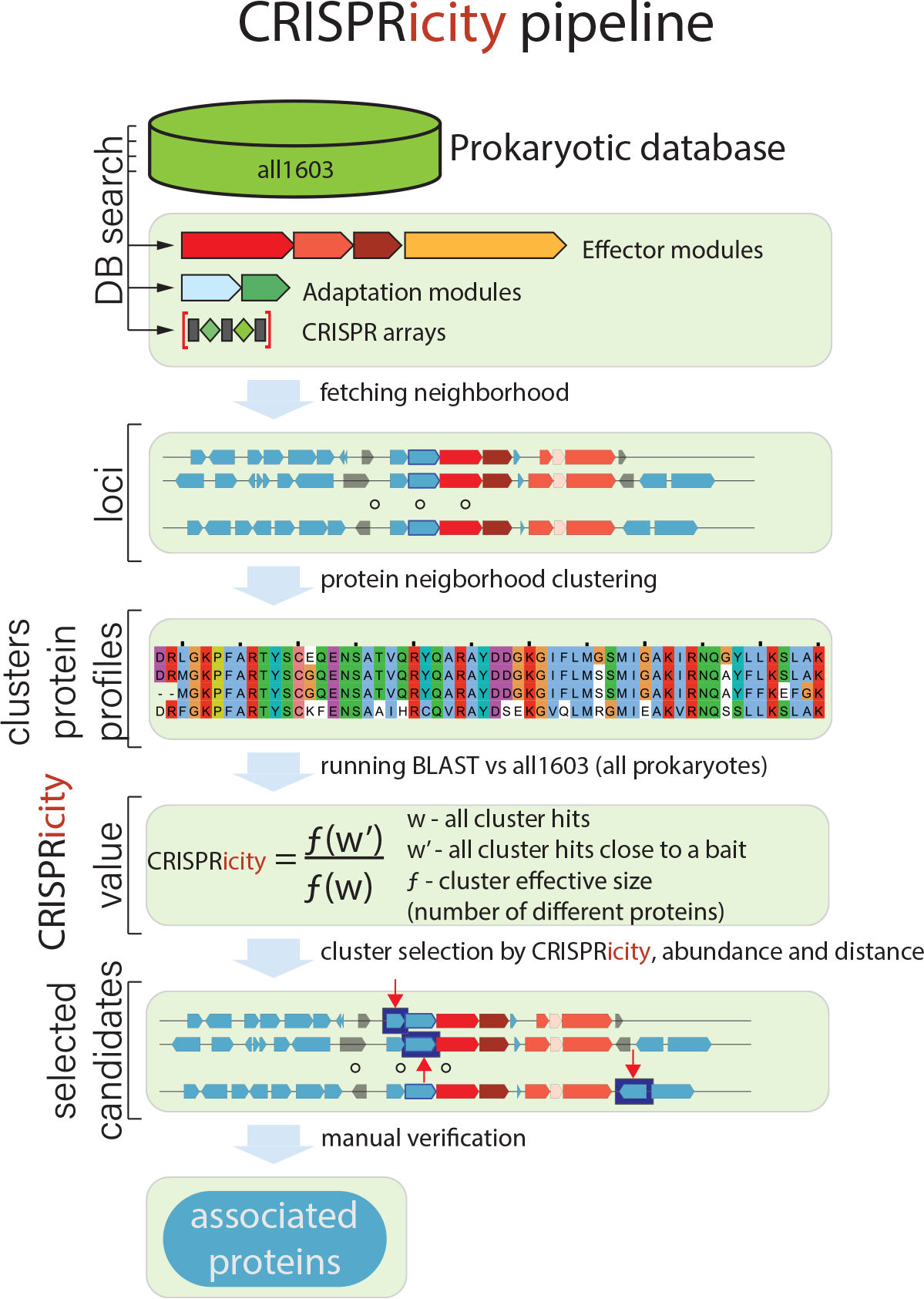
The computational pipeline for the analysis of the CRISPR-linked gene space.

Altogether, the above procedure yielded 468 putative distinct candidate CRISPR-linked genes (i.e. clusters of related sequences) (Figure 2A,B and Supporting Table S1). The 360 (predicted) clusters that included homologous proteins with sequences longer than 100 amino acids were thoroughly explored case by case using methods for sequence profile comparison and phylogenetic analysis (see Methods for details). From this initial list of candidates, 69 proteins were found to be previously unannotated and diverged variants of known Cas and accessory proteins; 79 were classified as likely previously undetected CRISPR-linked proteins; and the remaining 212 ones were inferred to be, most likely, irrelevant for the CRISPR function based on the genomic context analysis (are poorly conserved genes represented only in closely related genomes, common components of defense islands that only co-occur with *cas* genes, gene fragments, and open reading frames overlapping with CRISPR arrays; see Table S1). Previously missed members of Cas protein families known for their fast evolution, in particular, both the large and the small subunits of type I effector complexes and the type III small subunit, were identified in CRISPR-*cas* loci of most subtypes of both types (Figure 2B and Table S1 in the SI). Additionally, detection of previously unidentified Cas proteins was common for type IV systems that are typically carried by plasmids and appear to be the most diverged among the known CRISPR-Cas systems (Figure 2B and Table S1 in the SI). Strikingly, the distribution of the predicted accessory proteins among CRISPR-Cas variants was far more skewed than the distribution of previously undetected Cas proteins: the great majority of candidate accessory genes were detected in type III loci (Figure 2B and Table S1). When this manuscript was in the final stage of preparation, a preprint has been released describing the identification of 39 protein families associated with type III CRISPR-Cas systems (28); this list partially overlaps with list of CRISPR-linked genes detected in the present work. To assist future annotation of CRISPR-Cas loci we constructed new sequence alignments (profiles) for accessory proteins using the sequences detected in this study (Supporting Information Files 2 and 3).

**Figure 2.**
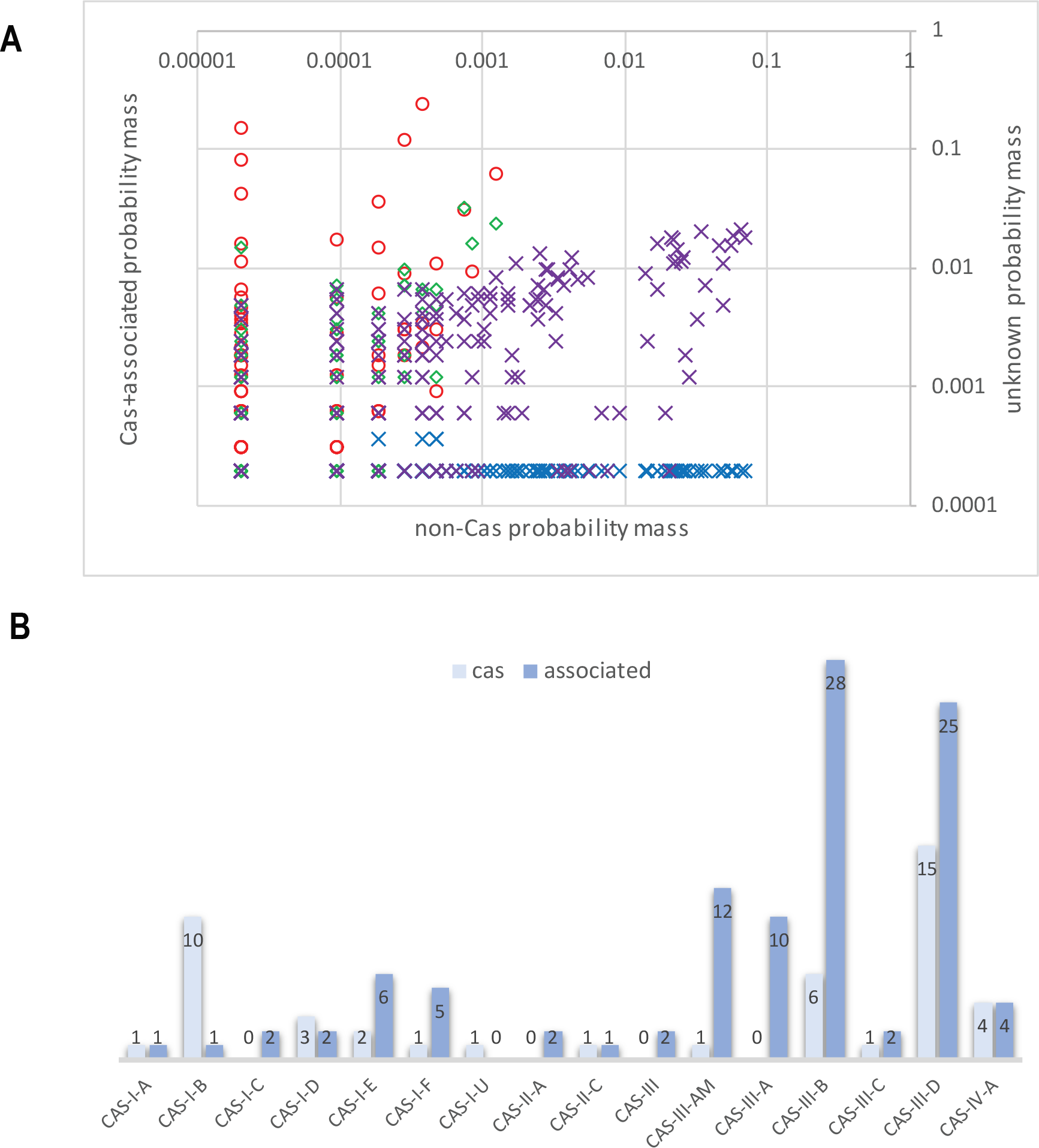
Protein clusters in CRISPR gene neighborhoods. A. Distribution of voxels in the CRISPRicity-abundance-distance space. Red circles: probability mass distribution for the union of *cas* and previously identified CRISPR-associated genes, voxels with CRISPR-index *I* > 2; blue crosses: probability mass distribution for the union of *cas* and previously identified CRISPR-associated genes, voxels with CRISPR-index *I* < 2; green diamonds: probability mass distribution for unknown genes, voxels with CRISPR-index *I* > 2 (candidate CRISPR-linked genes); purple crosses: probability mass distribution for unknown genes, voxels with CRISPR-index *I* < 2. B. Breakdown of previously undetected CRISPR-linked protein clusters by the CRISPR-Cas types and subtypes.

#### Predicted CRISPR-linked proteins: membrane connections and signal transduction

The set of predicted CRISPR-accessory proteins included those that have been shown to stably co-occur or even function jointly with specific CRISPR-Cas variants but have not been formally listed as CRISPR-linked. These included reverse transcriptase (RT) (17, 18), Argonaute family proteins (29, 30) and transposon-encoded proteins of the TniQ family (31). However, the majority of the predicted CRISPR-linked proteins have not been described previously. Below we discuss several examples of previously undetected CRISPR-linked genes that are representative of the findings made through the CRISPRicity analysis.

A major theme that is emerging from the case by case examination of the predicted CRISPR accessory genes is the potential membrane connection of CRISPR-Cas systems, especially, those of type III (Table S1). The most abundant of these genes is *corA* which encodes a widespread divalent cation channel in bacteria and archaea where it provides the primary route for electrophoretic Mg^2+^ uptake (32) (Figure 3A). A subset of CorA proteins are encoded in numerous subtype III-B CRISPR-*cas* loci. These loci show considerable diversity of genome architectures, and in many of them, the *corA* gene is adjacent to a gene coding for a DHH family nuclease (33) or even fused to it (eg. EDN71418.1 from *Beggiatoa sp*. PS). Moreover, some of these loci also contain a gene for another predicted nuclease (RNase), one of the NYN family (34) (Figure 3A). Structural analysis of CorA has shown that the protein is a pentamer, with each subunit contributing two transmembrane (TM) helices and containing a bulky cytosolic part (35, 36). The stable association of CorA with type III-B CRISPR-Cas strongly suggests a functional connection, and more specifically, a link between the CRISPR-mediated defense and membrane processes. Moreover, in a previous analysis, we have serendipitously detected evidence of recombination within III-B loci of Clostridium botulinum where *corA* segregated with the effector genes, suggestive of coordinated functions (37). However, predicting the CRISPR-linked functions of CorA more precisely is difficult. The nuclease connection suggests that CorA might regulate additional cleavage of the target and/or its transcripts DNA during the CRISPR response but more generally, it seems likely that CorA was exapted for membrane tethering of the respective CRISPR-Cas systems.

**Figure 3.**
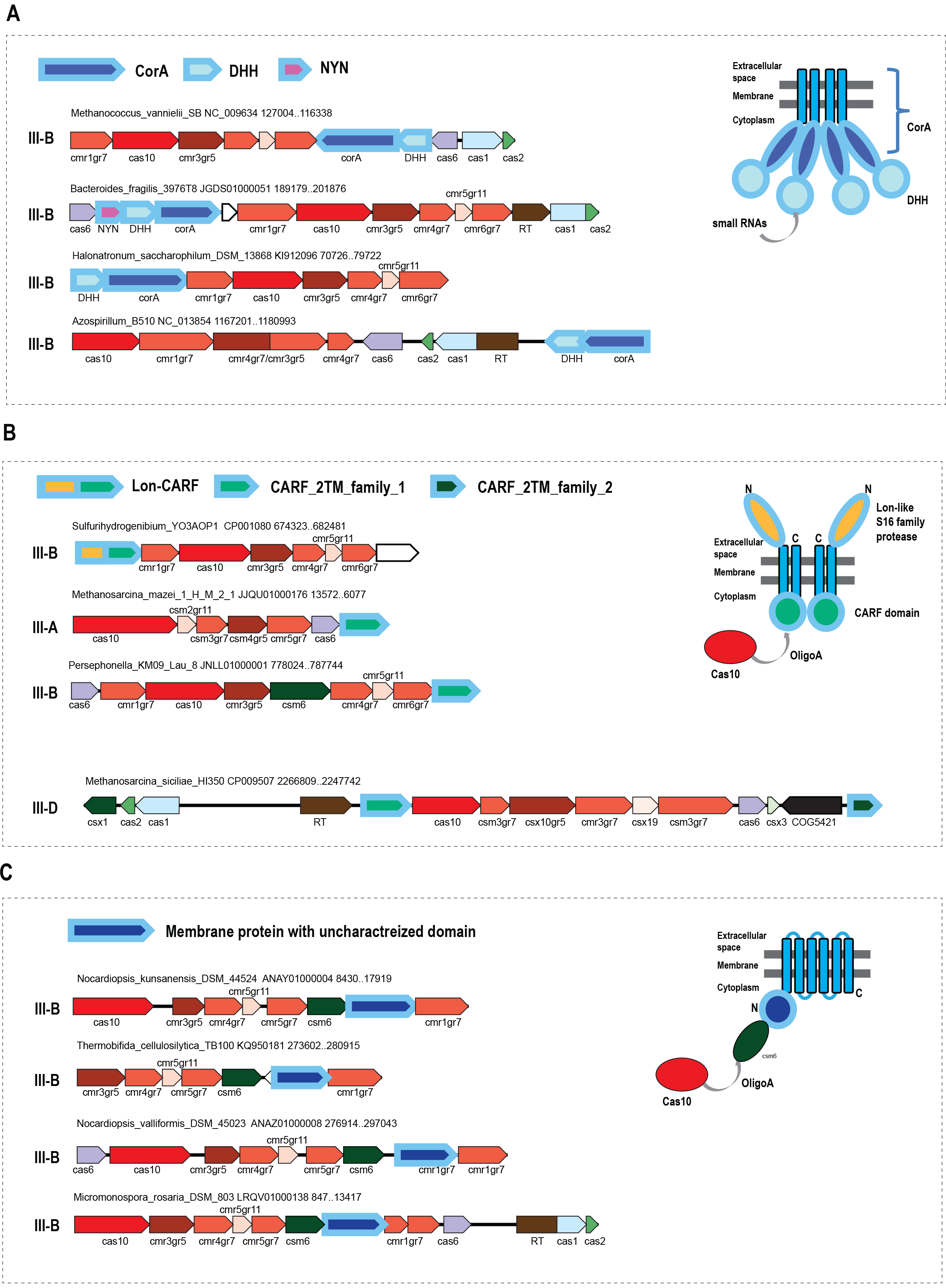
Locus organization of type III CRISPR-Cas systems containing predicted CRISPR-linked genes encoding membrane proteins. A. CorA, divalent cation membrane channel encoded in type III-B CRISPR-ca loci along with two distinct nucleases B. Membrane-associated CARF domain-containing proteins C. Uncharacterized membrane protein family in diverse type III loci For each locus, species name, genome accession number and the respective nucleotide coordinates and CRISPR-Cas system subtype are indicated. The genes in a representative locus are shown by block arrows which show the transcription direction. The scale of an arrow is roughly proportional to the respective gene length. Homologous genes and domains are color-coded, empty arrows show predicted genes without detectable homologs. On the right, models of the membrane topology of the predicted CRISPR-linked membrane proteins protein are shown according to the TMHMM predictions. Hypothetical interactions of the identified CRISPR-linked proteins with CRISPR-Cas system components are also depicted (see text). The *cas* gene names follow the current nomenclature (11); for several core *cas* genes, an extension specifies the gene group (gr5, gr6, gr7, groups 5, and 7 of the RAMP superfamily, respectively; gr8, large subunit of the effector complex; gr11, small subunit of the effector complex). Abbreviations and other gene names: RT, reverse transcriptase; DHH, DHH family nuclease; NYN, NYN family nuclease; Lon, Lon family protease; CARF, CRISPR-associated Rossmann fold domain; TM, transmembrane helix; COG5421, transposase of COG5421 family.

Numerous type III systems of subtypes A, B and D encompass genes that encode previously undetected, highly diverged proteins containing a CARF domain (38) and two predicted transmembrane helices; additionally, some of these proteins contain a fused, diverged Lon family (39) protease domain (Figure 3B). Membrane topology prediction suggests that the CARF domain faces the cytosol whereas the Lon protease is extracellular (Figure 3B). Given that the respective CRISPR-*cas* loci do not encode any other CARF domains but possess Cas10 proteins with predicted nucleotide polymerization activity, it appears most likely that the membrane-bound CARFs recognize signaling oligoA molecules synthesized by Cas10. However, the nature of the effector that could be activated by such binding remains unclear. The Lon family protease might fulfill this function in the case of the respective fusion proteins but we cannot currently predict the specific mechanism. In other cases, predicted membrane proteins are stably associated with type III CRISPR-Cas but contain no identifiable soluble domains that would provide for a specific functional prediction; nevertheless, it appears likely that such proteins anchor the CRISPR-Cas machinery in the bacterial membrane (Figure 3C).

A distinct variant of apparently degenerate I-E systems that lack Cas1, Cas2 and Cas3, and accordingly, cannot be active in either adaptation or target cleavage encode a predicted NTPase of the STAND superfamily (40) that, in addition to the P-loop NTPase domain, contains a cassette of TPR repeats (Figure 4A). The STAND NTPases that typically contain protein-protein interaction domains, such as TPR, are involved in various signal transduction networks that are poorly characterized in prokaryotes, but in eukaryotes, are involved in programmed cell death (40, 41). Phylogenetic analysis of STAND NTPases shows that the CRISPR-associated ones form a strongly supported branch (Figure 4A, Supporting Information File 4). Given the considerable diversity of gene arrangements in these loci, the monophyly of the NTPases implies their long-term association with CRISPR-Cas systems. Notably, in many cases, the gene adjacent to the NTPase gene encodes a predicted small membrane protein (Figure 4A) suggesting, once again, membrane association of CRISPR-Cas. This particular variant of subtype I-E is likely to perform a non-defense, probably, regulatory function and, through the STAND NTPase, might connect to signaling pathways that remain to be identified.

**Figure 4.**
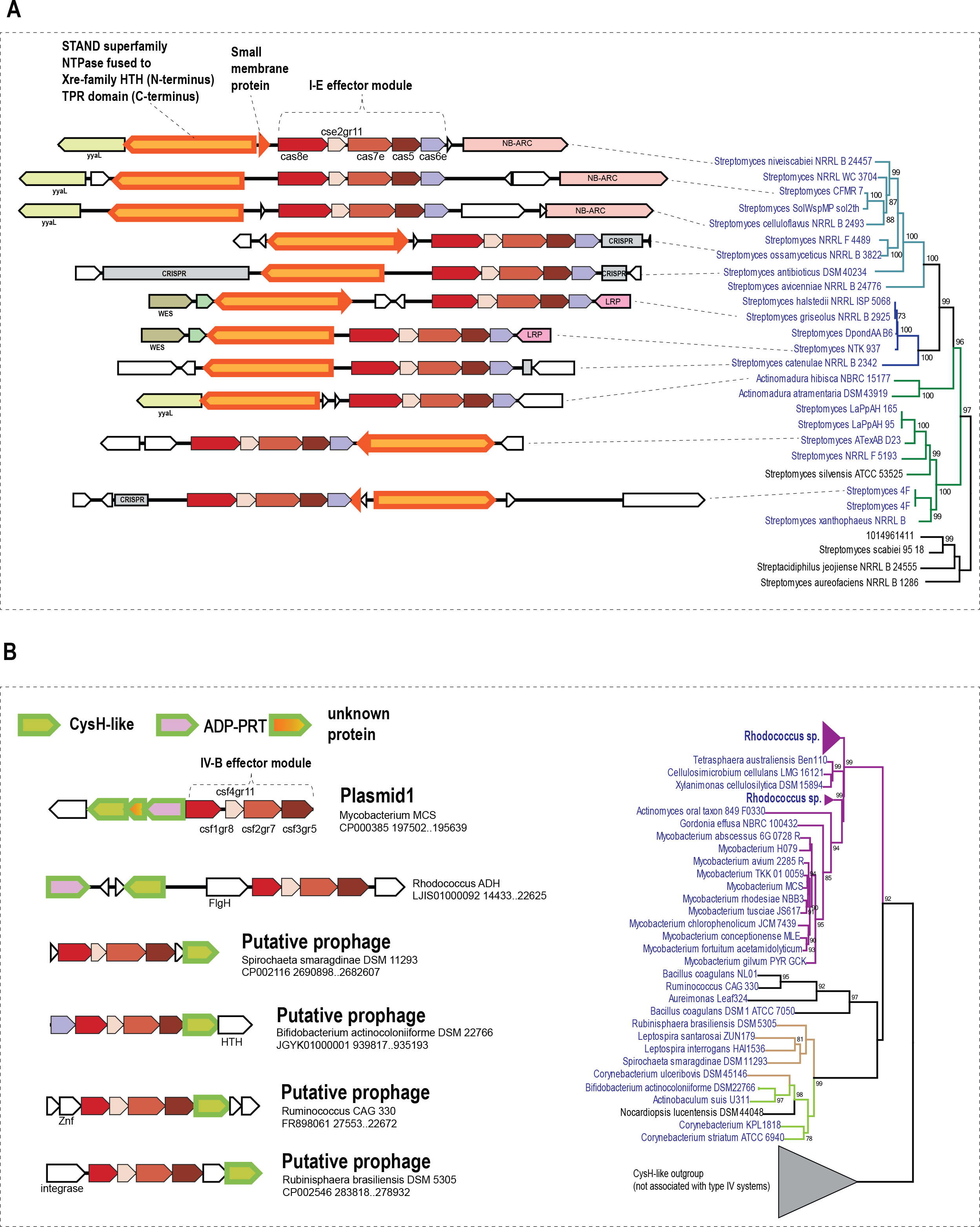
Locus organization of type I-E and type IV CRISPR-Cas systems containing predicted CRISPR-linked genes. A. STAND family NTPases encoded in minimal type I-E loci The clade of STAND NTPases associated with type I-E systems is shown on the right (complete tree is available at ftp://ftp.ncbi.nlm.nih.gov/pub/wolf/_suppl/CRISPRicity). Genomes in which the STAND NTPases gene is linked to a type I-E locus are shown in blue, and genomes in which there is no such link are shown in black. Colored branches denote three subfamilies (clusters) identified in this work. Support values greater than 70% are indicated for the respective branches. B. CysH family PAPS reductases encoded in type IV-B loci The clade of CysH family enzymes associated with type IV CRISPR-Cas systems is shown on the right (complete tree is available at ftp://ftp.ncbi.nlm.nih.gov/pub/wolf/_suppl/CRISPRicity). Genomes in which *cysH*-like genes are linked to type IV-B loci are shown in blue, and genomes in which there is no such link are shown in black. Colored branches denote three subfamilies (clusters) identified in this work. Support values greater than 70% are indicated for respective branches. The designations are as in Figure 3. CRISPR-arrays are shown by gray block arrows. Additional abbreviations and gene names: ADP-PRT, ADP phosphoribosyltransferase; LRP, LRP family transcriptional regulator; NB_ARC, STAND NTPase fused to TPR-repeats (distinct from the predicted CRISPR-linked STAND NTPase); HTH, helix-turn-helix DNA-binding domain; FlhG, MinD-like ATPase involved in chromosome partitioning or flagellar assembly; SSB, single-stranded DNA-binding protein; N6-MTase, N6 adenosine methylase.

Many subtype IV-B loci that are typically located on plasmids or predicted prophages (Figure 4B) encode a predicted enzyme of the CysH family which belongs to the adenosine 5′-phosphosulfate (PAPS) reductase family (42). Some of these loci also encode a predicted enzyme of the ADP-ribosyltransferase family (43) (Figure 4B). Similarly to the subtype I-E systems discussed above as well as “minimal” I-F systems encoded by Tn7-like transposons (31), type IV systems lack nucleases that could cleave the target DNA, and therefore, can be predicted to perform non-defense functions similarly to transposon-encoded CRISPR-Cas systems (31). Analogously to the case of the STAND NTPases, the CRISPR-associated CysH homologs comprise a well-supported clade in the phylogenetic tree of the CysH protein family (Figure 4B, Supporting Information File 5). As with other predicted CRISPR accessory genes, the CysH-like enzyme and the associated proteins might play a role in a signal transduction pathway connecting CRISPR-Cas with cellular regulatory networks and perhaps stabilizing the prophages and plasmids in the host bacteria.

#### Coevolution of predicted CRISPR-linked genes with bona fide cas genes

The predicted CRISPR-linked genes recur in multiple CRISPR-*cas* loci and accordingly can be predicted to contribute to the functions of the respective CRISPR-Cas systems. We sought to explore the linkage between these genes and CRISPR-Cas at a deeper level. To this end, we performed an analysis of potential coevolution between those of the predicted accessory genes that are sufficiently widespread and well-conserved with signature *cas* genes. Phylogenetic trees were constructed for the analyzed CRISPR-linked genes and compared to trees of *cas* genes and, as a control, to those for 16S RNA (a proxy for the species tree of the respective microbes), and the evolutionary distances between the respective genes from different organisms extracted from the trees were compared (see Methods for details). Notably, the evolutionary distances for the *corA* genes were strongly, positively correlated with the distances for *cas10* whereas none of these genes showed comparative correlation with 16S RNA (Figure 5A-C, Supporting Information Files 6-10) suggesting that the genes within the CRISPR-*cas* loci including *corA* coevolve but the loci themselves spread largely via horizontal transfer. In the case of the membrane-associated CARF-domain proteins, highly significant correlation was observed between the trees for these proteins and both Cas10 and 16S RNA (Figure 5 D-F) suggesting that the evolution of the respective subset of type III CRISPR-Cas systems, including the accessory proteins predicted here, involved a major vertical component. In contrast, in the case of RT associated with type III systems, the correlations between the RT trees and those for both cas10 and 16 S RNA were relatively weak (Figure 5 G-I), in agreement with the previous conclusion that the RT-containing adaptation modules largely behaved as distinct evolutionary units (17). Thus, comparative analysis of phylogenetic trees highlights distinct patterns of evolution among CRISPR-Cas systems but on the whole, presents strong evidence of coevolution and implies tight functional association between (at least) the most common of the predicted CRISPR accessory genes and the effector modules of the respective systems.

**Figure 5.**
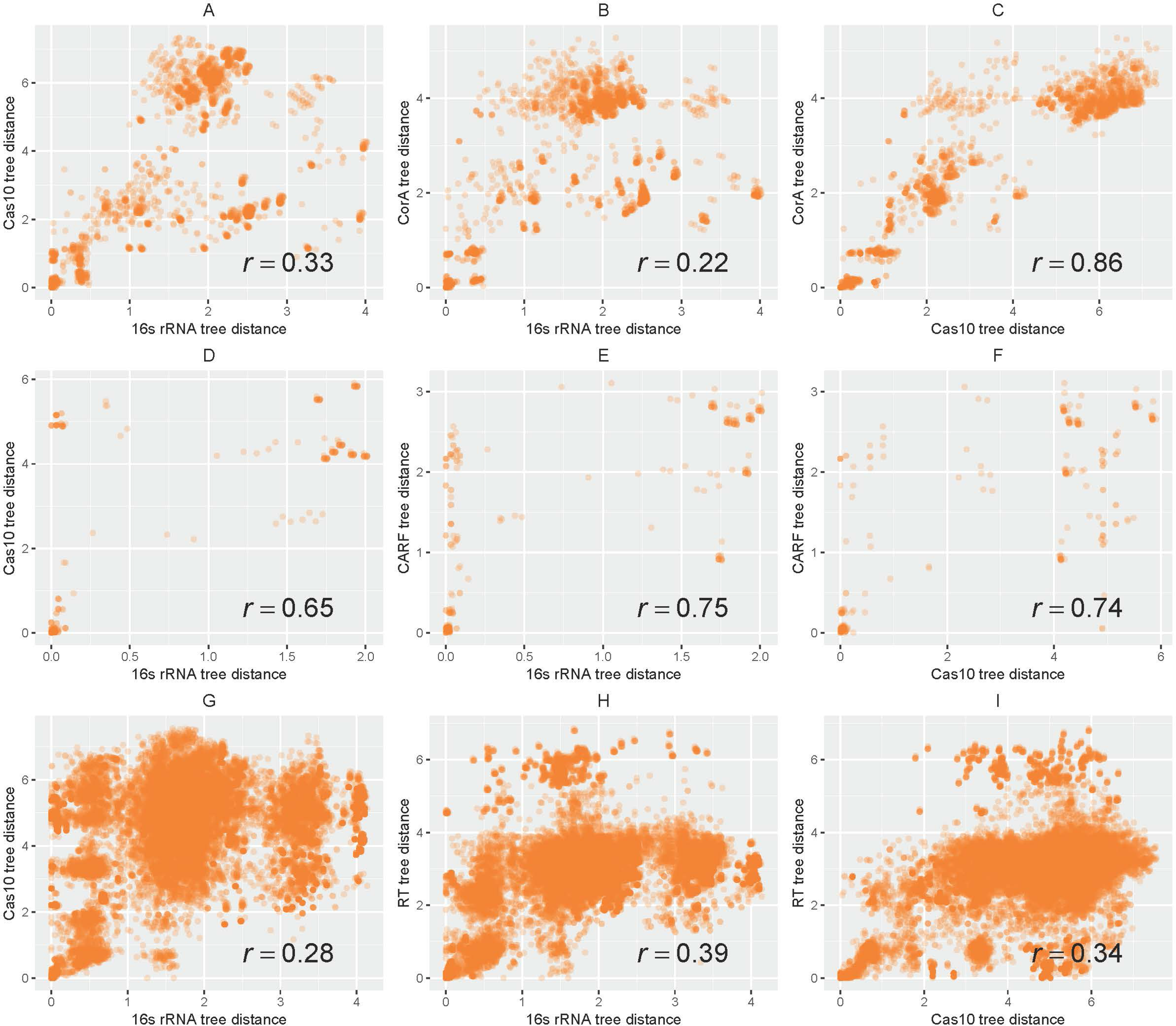
Coevolution of predicted CRISPR-linked genes with signature *cas* genes. The panels show plots of pairwise distances between predicted CRISPR-linked gene products (CorA, membrane-associated CARF-domain proteins and RT), Cas10 and 16S rRNA estimated from the respective phylogenetic trees. The Spearman rank correlation coefficient is indicated on each plot.

## CONCLUSIONS

We developed a computational strategy to predict genes that are functionally linked to CRISPR-Cas systems. Exhaustive case by case analysis of the detected CRISPR-linked genes shows that, despite the absence of rigorous statistical framework, this “CRISPRicity” strategy yields sets of genes that are highly enriched in confident predictions of functional association. Clearly, this approach can be readily generalized beyond CRISPR-Cas, to systematically explore functional links for any other systems encoded in microbial genomes. In biological terms, the CRISPRicity analysis reveals remarkable functional complexity of type III CRISPR-Cas systems that seems to substantially exceed that other CRISPR-Cas types. Major themes among the previously unnoticed CRISPR-linked genes identified here are the predicted membrane association and connections to signal transduction pathways. These findings imply that entire layers of CRISPR-Cas biology remain unexplored and open up many experimental directions.

## MATERIALS AND METHODS

### Prokaryotic Genome Database

Prokaryotic database that consisted of 4,961 completely assembled genomes and 43,599 partial genomes, or 6,342,452 nucleotide sequences altogether (genome partitions, such as chromosomes and plasmids, and contigs), was assembled from archaeal and bacterial genomic sequences downloaded from the NCBIFTP database (ftp://ftp.ncbi.nlm.nih.gov/genomes/all/) in March 2016. Default open reading frame (ORF) annotation available on the FTP site was used for well annotated genomes (coding density > 0.6 coding sequences per kilobase), and the rest of the genomes were annotated with Meta-GeneMark (44) using the standard model MetaGeneMark_v1.mod (Heuristic model for genetic code 11 and GC 30).

### CRISPR array detection and annotation

The CRISPR arrays were identified as previously described (45). Briefly, the Prokaryotic Genome Database was scanned with CRISPRFinder (46) and PILER-CR (47) using default parameters. The search identified 61,581 and 49,817 CRISPR arrays respectively. The union of the search results with the two methods were taken as the set of 65,194 predicted CRISPR arrays; the CRISPRFinder prediction was accepted in cases of overlap. To eliminate spurious CRISPR array predictions, arrays of unknown type that did not produce reliable BLASTN hits (90% identity and 90% coverage) into CRISPR arrays of known type were discarded. This filtering resulted in 42,352 CRISPR arrays that were taken as the final prediction for the subsequent analyses.

### Detection and annotation of CRISPR-Cas proteins

The translated prokaryotic database was searched with PSI-BLAST (48) using the previously described CRISPR-Cas protein profiles (11, 49, 50) with an e-value cutoff of 10^−4^ and effective database size set to 2×10^7^.

### Bait islands

For the purpose of identification of previously undetected CRISPR-linked genes, three groups of “baits” were selected:

1) All CRISPR arrays from the final set of predictions
2) Effector modules, i.e. all interference-related genes detected using Cas protein profiles
3) Adaptation modules, i.e. *cas1* genes located within 10 kb of a *cas2* and/or *cas4* gene (this additional criterion was adopted because of the existence of non-CRISPR-associated *cas1* homologs).

Bait islands, a data structure describing genomic neighborhoods of the above 3 classes of baits, were constructed by annotating all ORFs within 10 kb upstream and downstream of the baits. The ORFs were annotated using 30,953 COG, pfam, and cd protein profiles from the NCBI CDD database (51) and 217 custom CRISPR-Cas profiles (49) using PSI-BLAST profile search with the same parameters as above.

### Construction of protein clusters

Clusters of homologous proteins were constructed for all ORFs detected in the bait islands using the following iterative procedure:

1. all proteins were clustered using UCLUST (52) with the sequence similarity threshold of 0.3 forming permissive cluster set;
2. for each permissive cluster with 3 or more proteins, its members were clustered using UCLUST with sequence similarity threshold of 0.9, forming strict clusters inside the permissive cluster. The number of strict clusters within a permissive cluster was taken to represent the effective number of sequences in the given permissive cluster;
3. for each permissive cluster, representatives of all constituent strict clusters were aligned using MAFFT (53). The resulting alignments were used as queries to initiate a PSI-BLAST search against all sequences within the permissive cluster. Sequences that did not produce a significant hit (e-value cutoff of 10^—4^) were removed. This step was repeated until convergence;
4. three iterations of steps 2 and 3 were repeated with the updated set of the permissive clusters;
5. the final clustering step using UCLUST with sequence similarity threshold of 0.5 was performed with all remaining singleton sequences;

The 16,433 clusters with effective size of 3 and greater that were constructed using this procedure were used for further analysis.

### Measures of CRISPR-Cas association

Alignments of the permissive clusters was used to initiate a PSI-BLAST search against the prokaryotic sequence database with an e-value cutoff of 10^−4^ and effective database size set to 2×10^7^. All hits covering less than 40% of the query profile were discarded. When multiple queries produced overlapping hits (using the overlap threshold of 25%) to the same target sequence, the corresponding target segment was assigned to the highest-scoring query.

All target segments assigned to the same query were, once again, clustered using UCLUST with the sequence similarity threshold of 0.3. Clusters that did not contain any sequence from the query profiles were removed.

For the sets of hits assigned to a particular query profile, the effective number of sequences was calculated as described above for:

1. all sequences retrieved from the Prokaryotic Genome Database retrieved by the given profile;
2. sequences from all bait islands
3. sequences from the CRISPR array islands
4. sequences from the effector module islands
5. sequences from the adaptation module islands

The aggregate measure of CRISPR association (CRISPRicity) was calculated for each permissive cluster as the ratio of the effective number of sequences in the bait islands to the effective number of sequences in the entire database that were associated with the given cluster.

Each gene in a specific bait island can be characterized by its genomic distance from the respective bait, i.e. the number of gene between the given gene and the bait (genes directly adjacent to the bait and the baits themselves were assigned the distance of 0). The median of the distance to the closest bait across all representatives was used to characterize the permissive cluster.

### Selection of candidate CRISPR-linked protein clusters

The CRISPRicity-abundance-distance space, embedding all permissive clusters, was partitioned into 1,000 voxels (volume elements) as follows. The CRISPRicity range (from 0 to 1) was split into 10 equal intervals. The abundance (the effective number of sequences in bait islands) range was split into 10 intervals in log space with a step of 0.3 decimal log units (factor of ~2.0), starting from 1; all clusters with abundance of 502 = 10^0.3×9^ or greater were assigned to the last interval. The distance range was split into 10 intervals with a step of 1, starting from the distance of 0 (adjacent to the bait); clusters with distances of 9 and above were assigned to the last interval. This 10×10×10 grid formed the 1,000 voxels. Within each voxel, the known Cas (together with Cas-associated) clusters and non-Cas clusters were counted; their probability masses (*D_c_* and *D_n_* respectively) were calculated as the counts divided by the total number of such genes. Voxels with the CRISPR-index (ratios of densities) *I* = *D_c_*/*D_n_* > 2 were selected for further analysis.

### Phylogenetic analysis

The 16s rRNA tree was constructed for all organisms from the prokaryotic database where both 16S SSU rRNA and CRISPR-Cas Type III system were found. The 16S rRNA genes were identified using BLASTN (48) search with the *Pyrococcus sp*. NA2 16S rRNA as the query for archaeal genomes and the *Escherichia coli* K-12 ER3413 16S rRNA as a query for bacterial genomes (word size of 8 and dust filtering off). The best scoring BLASTN hits with at least 80% coverage and 70% identity were taken for each organism, 16S rRNA sequences were aligned using MAFFT (53), and a phylogenetic tree was constructed using FastTree (54) with gamma-distributed site rates and GTR evolutionary model. Protein sequences were iteratively aligned using hhalign (55) starting from MUSCLE (56) alignments of UCLUST clusters (similarity cutoff of 0.5). Approximate ML trees were built from these alignments using FastTree with gamma-distributed site rates and WAG evolutionary model.

### Case by case analysis and annotation of permissive clusters of putative CRISPR-linked proteins

PSI-BLAST searches with COG, pfam, and cd protein profiles from NCBI CDD database (51) and with the custom CRISPR-Cas profiles (49) were run against a database made from consensus sequences (57) of the permissive clusters with an e-value cutoff of 10^−4^ and effective database size set to 2×10^7^.

Iterative profile searches using PSI-BLAST (48), with a cut-off e-value of 0.01, and composition based-statistics and low complexity filtering turned off, were used to search for distantly similar sequences in NCBI’s non-redundant (NR) database. Another sensitive method for remote sequence similarity detection, HHpred, was used with default parameters (55). Additionally, clusters were annotated using HHSearch (55) comparison between cluster-derived HMM profiles and CDD-derived HMM profiles. The results with an HHSearch probability score greater than 80% were recorded.

Protein secondary structure was predicted using Jpred (58). Transmembrane segments were predicted using the TMMHMM v. 2.0c program with default parameters (59).

A fraction of the permissive protein clusters was found to comprise CRISPR arrays falsely annotated as protein-coding sequences. Clusters containing more than 10% of sequences overlapping with known CRISPR arrays or matching CRISPR repeats with 90% identity and 90% coverage in BLASTN search were identified and discarded from further analysis.

## Acknowledgements

We thank Koonin group members for useful discussions.

## SUPPORTING INFORMATION

Table S1. Analysis of candidate CRISPR-linked protein clusters

### SUPPORTING INFORMATION FILES (available via ftp)

**Additional Information File 1. Genome neighborhoods of all baits (CRISPR arrays, effector genes and adaptation genes)**

Islands.tar.gz at ftp://ftp.ncbi.nlm.nih.gov/pub/wolf/_suppl/CRISPRicity

**Additional Information File 2. Multiple sequence alignments (profiles) for CRISPR-Cas accessory genes**

NewProfiles.tar.gz at ftp://ftp.ncbi.nlm.nih.gov/pub/wolf/_suppl/CRISPRicity

**Additional Information File 3. List of sequence alignments (profiles) for CRISPR-Cas accessory genes**

icity-profiles.xlsx at ftp://ftp.ncbi.nlm.nih.gov/pub/wolf/_suppl/CRISPRicity

**Additional Information File 4. Phylogenetic analysis of STAND NTPase family**

STAND.tar.gz at ftp://ftp.ncbi.nlm.nih.gov/pub/wolf/_suppl/CRISPRicity (tree in Newick format, sequences in FASTA format)

**Additional Information File 5. Phylogenetic analysis of CysH family**

CysH.tar.gz at ftp://ftp.ncbi.nlm.nih.gov/pub/wolf/_suppl/CRISPRicity (tree in Newick format, sequences in FASTA format)

**Additional Information File 6. Phylogenetic analysis of CRISPR-Cas associated RTs**

RT.tar.gz at ftp://ftp.ncbi.nlm.nih.gov/pub/wolf/_suppl/CRISPRicity (tree in Newick format, sequences in FASTA format)

**Additional Information File 7. Phylogenetic analysis of CARF family**

CARF.tar.gz at ftp://ftp.ncbi.nlm.nih.gov/pub/wolf/_suppl/CRISPRicity (tree in Newick format, sequences in FASTA format)

**Additional Information File 8. Phylogenetic analysis of CorA family**

CorA.tar.gz at ftp://ftp.ncbi.nlm.nih.gov/pub/wolf/_suppl/CRISPRicity (tree in Newick format, sequences in FASTA format)

**Additional Information File 9. Phylogenetic analysis of Cas10 family**

Cas10.tar.gz at ftp://ftp.ncbi.nlm.nih.gov/pub/wolf/_suppl/CRISPRicity (tree in Newick format, sequences in FASTA format)

**Additional Information File 10. Phylogenetic analysis of 16S rRNA**

16sRRNA.tar.gz at ftp://ftp.ncbi.nlm.nih.gov/pub/wolf/_suppl/CRISPRicity (tree in Newick format, sequences in FASTA format)

## References

1. Sorek, R, Lawrence, CM, & Wiedenheft, B (2013) CRISPR-mediated adaptive immune systems in bacteria and archaea. Annu Rev Biochem 82:237–266

2. Wright, AV, Nunez, JK, & Doudna, JA (2016) Biology and Applications of CRISPR Systems: Harnessing Nature’s Toolbox for Genome Engineering. Cell 164(1-2):29–44

3. Komor, AC, Badran, AH, & Liu, DR (2016) CRISPR-Based Technologies for the Manipulation of Eukaryotic Genomes. Cell http://dx.doi.org/10.1016/j.cell.2016.10.044

4. Mohanraju, P, et al. (2016) Diverse evolutionary roots and mechanistic variations of the CRISPR-Cas systems. Science 353(6299):aad5147

5. Barrangou, R & Horvath, P (2017) A decade of discovery: CRISPR functions and applications. Nat Microbiol 2:17092

6. Amitai, G & Sorek, R (2016) CRISPR-Cas adaptation: insights into the mechanism of action. Nat Rev Microbiol 14(2):67–76

7. Sternberg, SH, Richter, H, Charpentier, E, & Qimron, U (2016) Adaptation in CRISPR-Cas Systems. Mol Cell 61(6):797–808

8. Charpentier, E, Richter, H, van der Oost J, & White MF (2015) Biogenesis pathways of RNA guides in archaeal and bacterial CRISPR-Cas adaptive immunity. FEMS Microbiol Rev 39(3):428–441

9. Plagens, A, Richter, H, Charpentier, E, & Randau, L (2015) DNA and RNA interference mechanisms by CRISPR-Cas surveillance complexes. FEMS Microbiol Rev 39(3):442–463

10. Nishimasu, H & Nureki, O (2017) Structures and mechanisms of CRISPR RNA-guided effector nucleases. Curr Opin Struct Biol 43:68–78

11. Makarova, KS, et al. (2015) An updated evolutionary classification of CRISPR-Cas systems. Nat Rev Microbiol 13(11):722–736

12. Koonin, EV, Makarova, KS, & Zhang F (2017) Diversity, classification and evolution of CRISPR-Cas systems. Curr Opin Microbiol 37:67–78

13. Doudna, JA & Charpentier, E (2014) Genome editing. The new frontier of genome engineering with CRISPR-Cas9. Science 346(6213):1258096

14. van der Oost J, Westra, ER, Jackson, RN, & Wiedenheft, B (2014) Unravelling the structural and mechanistic basis of CRISPR-Cas systems. Nat Rev Microbiol 12(7):479–492

15. Jiang, F & Doudna, JA (2015) The structural biology of CRISPR-Cas systems. Curr Opin Struct Biol 30:100–111

16. Makarova, KS, Wolf, YI, & Koonin, EV (2013) The basic building blocks and evolution of CRISPR-cas systems. Biochem Soc Trans 41(6):1392–1400

17. Silas, S, et al. (2017) On the Origin of Reverse Transcriptase-Using CRISPR-Cas Systems and Their Hyperdiverse, Enigmatic Spacer Repertoires. MBio 8(4)

18. Silas, S, et al. (2016) Direct CRISPR spacer acquisition from RNA by a natural reverse transcriptase-Cas1 fusion protein. Science 351(6276):aad4234

19. Anantharaman, V, Makarova, KS, Burroughs, AM, Koonin, EV, & Aravind, L (2013) Comprehensive analysis of the HEPN superfamily: identification of novel roles in intra-genomic conflicts, defense, pathogenesis and RNA processing. Biol Direct 8:15

20. Kazlauskiene, M, Kostiuk, G, Venclovas, C, Tamulaitis, G, & Siksnys, V (2017) A cyclic oligonucleotide signaling pathway in type III CRISPR-Cas systems. Science 357(6351):605–609

21. Niewoehner, O, et al. (2017) Type III CRISPR-Cas systems produce cyclic oligoadenylate second messengers. Nature

22. Makarova, KS, Wolf, YI, Snir, S, & Koonin, EV (2011) Defense islands in bacterial and archaeal genomes and prediction of novel defense systems. J Bacteriol 193(21):6039–6056

23. Jacob, F & Monod, J (1961) Genetic regulatory mechanisms in the synthesis of proteins. J. Mol. Biol. 3:318–356

24. Wolf, YI, Rogozin, IB, Kondrashov, AS, & Koonin, EV (2001) Genome alignment, evolution of prokaryotic genome organization and prediction of gene function using genomic context. Genome Res. 11:356–372

25. Beckwith, J (2011) The operon as paradigm: normal science and the beginning of biological complexity. J Mol Biol 409(1):7–13

26. Rogozin, IB, et al. (2002) Connected gene neighborhoods in prokaryotic genomes. Nucleic Acids Res 30(10):2212–2223

27. Makarova, KS, Aravind, L, Grishin, NV, Rogozin, IB, & Koonin, EV (2002) A DNA repair system specific for thermophilic Archaea and bacteria predicted by genomic context analysis. Nucleic Acids Res 30(2):482–496

28. Shah SAA, O. S., Behler, J., Han, W., She, Q., Hess, W. R., Garrett, R. A., Backofen, R. (2018) Conserved accessory proteins encoded with archaeal and bacterial Type III CRISPR-Cas gene cassettes that may specifically modulate, complement or extend interference activity. https://www.biorxiv.org/content/early/2018/02/09/262675

29. Swarts, DC, et al. (2014) The evolutionary journey of Argonaute proteins. Nat Struct Mol Biol 21(9):743–753

30. Kaya, E, et al. (2016) A bacterial Argonaute with noncanonical guide RNA specificity. Proc Natl Acad Sci U S A 113(15):4057–4062

31. Peters, JE, Makarova, KS, Shmakov, S, & Koonin, EV (2017) Recruitment of CRISPR-Cas systems by Tn7-like transposons Proc Natl Acad Sci U S A pii: 201709035. doi: 10.1073/pnas.1709035114

32. Maguire, ME (2006) The structure of CorA: a Mg(2+)-selective channel. Curr Opin Struct Biol 16(4):432–438

33. Aravind, L & Koonin, EV (1998) A novel family of predicted phosphoesterases includes Drosophila prune protein and bacterial RecJ exonuclease. Trends Biochem Sci 23(1):17–19

34. Anantharaman, V & Aravind L (2006) The NYN domains: novel predicted RNAses with a PIN domain-like fold. RNA Biol 3(1):18–27

35. Matthies, D, et al. (2016) Cryo-EM Structures of the Magnesium Channel CorA Reveal Symmetry Break upon Gating. Cell 164(4):747–756

36. Lerche, M, Sandhu, H, Flockner, L, Hogbom, M, & Rapp, M (2017) Structure and Cooperativity of the Cytosolic Domain of the CorA Mg(2+) Channel from Escherichia coli. Structure 25(8):1175–1186 e1174

37. Puigbo, P, Makarova, KS, Kristensen, DM, Wolf, YI, & Koonin, EV (2017) Reconstruction of the evolution of microbial defense systems. BMC Evol Biol 17(1):94

38. Makarova, KS, Anantharaman, V, Grishin, NV, Koonin, EV, & Aravind, L (2014) CARF and WYL domains: ligand-binding regulators of prokaryotic defense systems. Front Genet 5:102

39. Smith, CK, Baker, TA, & Sauer, RT (1999) Lon and Clp family proteases and chaperones share homologous substrate-recognition domains. Proc Natl Acad Sci U S A 96(12):6678–6682

40. Leipe, DD, Koonin, EV, & Aravind, L (2004) STAND, a class of P-loop NTPases including animal and plant regulators of programmed cell death: multiple, complex domain architectures, unusual phyletic patterns, and evolution by horizontal gene transfer. J Mol Biol 343(1):1–28

41. Koonin, EV & Aravind, L (2002) Origin and evolution of eukaryotic apoptosis: the bacterial connection. Cell Death Differ 9(4):394–404

42. Bick, JA, Dennis, JJ, Zylstra, GJ, Nowack, J, & Leustek, T (2000) Identification of a new class of 5′-adenylylsulfate (APS) reductases from sulfate-assimilating bacteria. J Bacteriol 182(1):135–142

43. Jeong, BR, et al. (2011) Structure function analysis of an ADP-ribosyltransferase type III effector and its RNA-binding target in plant immunity. J Biol Chem 286(50):43272–43281

44. Besemer, J, Lomsadze, A, & Borodovsky, M (2001) GeneMarkS: a self-training method for prediction of gene starts in microbial genomes. Implications for finding sequence motifs in regulatory regions. Nucleic Acids Res 29(12):2607–2618

45. Shmakov, SA, et al. (2017) The CRISPR Spacer Space Is Dominated by Sequences from Species-Specific Mobilomes. MBio 8(5)

46. Grissa I, Vergnaud, G, & Pourcel, C (2007) CRISPRFinder: a web tool to identify clustered regularly interspaced short palindromic repeats. Nucleic Acids Res 35(Web Server issue):W52–57

47. Edgar, RC (2007) PILER-CR: fast and accurate identification of CRISPR repeats. BMC Bioinformatics 8:18

48. Altschul, SF, et al. (1997) Gapped BLAST and PSI-BLAST: a new generation of protein database search programs. Nucleic Acids Res 25(17):3389–3402

49. Makarova, KS & Koonin, EV (2015) Annotation and Classification of CRISPR-Cas Systems. Methods Mol Biol 1311:47–75

50. Shmakov, S, et al. (2017) Diversity and evolution of class 2 CRISPR-Cas systems. Nat Rev Microbiol

51. Marchler-Bauer A, et al. (2015) CDD: NCBI’s conserved domain database. Nucleic Acids Res 43(Database issue):D222–226

52. Edgar, RC (2010) Search and clustering orders of magnitude faster than BLAST. Bioinformatics 26(19):2460–2461

53. Katoh, K & Standley DM (2013) MAFFT multiple sequence alignment software version 7: improvements in performance and usability. Mol Biol Evol 30(4):772–780

54. Price, MN, Dehal, PS, & Arkin, AP (2010) FastTree 2––approximately maximum-likelihood trees for large alignments. PLoS One 5(3):e9490

55. Soding, J (2005) Protein homology detection by HMM-HMM comparison. Bioinformatics 21(7):951–960

56. Edgar, RC (2004) MUSCLE: multiple sequence alignment with high accuracy and high throughput. Nucleic Acids Res 32(5):1792–1797

57. Yutin, N, Makarova, KS, Mekhedov, SL, Wolf, YI, & Koonin, EV (2008) The deep archaeal roots of eukaryotes. Mol Biol Evol 25(8):1619–1630

58. Drozdetskiy, A, Cole, C, Procter, J, & Barton, GJ (2015) JPred4: a protein secondary structure prediction server. Nucleic Acids Res 43(W1):W389–394

59. Krogh, A, Larsson, B, von Heijne G, & Sonnhammer, EL (2001) Predicting transmembrane protein topology with a hidden Markov model: application to complete genomes. J Mol Biol 305(3):567–580

